# Diversity of feature selectivity in macaque visual cortex arising from limited number of broadly-tuned input channels

**DOI:** 10.1101/291229

**Authors:** Yamni S. Mohan, Jaikishan Jayakumar, Errol K.J. Lloyd, Ekaterina Levichkina, Trichur R. Vidyasagar

## Abstract

Spikes (action potential) responses of most primary visual cortical cells in the macaque are sharply tuned for the orientation of a line or an edge and neurons preferring similar orientations are clustered together in cortical columns. The preferred stimulus orientation of these columns span the full range of orientations, as observed in recordings of spikes, which represent the outputs of cortical neurons. However, when we imaged also the thalamic input to these cells that occur on a larger spatial scale, we found that the orientation domain map of the primary visual cortex did not show the diversity of orientations exhibited by signals representing outputs of the cells. This map was dominated by just the one orientation that is most commonly represented in subcortical responses. This supports cortical feature selectivity and columnar architecture being built upon feed-forward signals transmitted from the thalamus in a very limited number of broadly-tuned input channels.

---

The extracellularly recorded spike responses of most neurons in the primary visual cortex (V1) show a preference for a narrow range of orientations of lines and edges. Seen on a broader scale, cortical area V1 also shows a clustering of cells tuned to similar preferred orientations in the form of orientation columns (Hubel and Wiesel, 1968). Not only is the neural basis for such orientation selectivity still highly debated (Priebe and Ferster, 2012; Vidyasagar and Eysel, 2015), but no scheme that explains *both* orientation selectivity and the architecture of orientation columns emerging from a single thalamocortical connectivity pattern has been experimentally demonstrated. The clue for such a unifying scheme may lie in two major findings. One is the presence of a bias for stimulus orientation seen in responses of cells in the retina and lateral geniculate nucleus (LGN), now demonstrated in every species studied so far (Levick and Thibos, 1982; Vidyasagar and Urbas, 1982; Shou and Leventhal, 1989; Passaglia et al., 2002; Xu et al., 2002; Tan et al., 2011; Van Hooser et al., 2013; Sun et al., 2016; Antinucci and Hindges, 2018). The other is the finding that the stimulus preference of these subcortical cells is not distributed evenly for all possible orientations, but the preference peaks for just a few orientations (Levick and Thibos, 1982; Vidyasagar and Urbas, 1982; Shou and Leventhal, 1989; Passaglia et al., 2002). It has been proposed, that as in the case of trichromatic colour vision, with its three broadly-tuned cone types forming the basis for the whole range of cortical hue preferences, the full gamut of orientation preferences seen cortically may be derived from a limited number of broadly tuned channels in its input (Vidyasagar, 1985: Vidyasagar and Eysel, 2015).

We tested this proposal using a novel method of analysing data from optical imaging of intrinsic signals (OI) evoked in the macaque V1 in response to drifting sinusoidal gratings of different orientations. The bulk of the haemodynamic signals imaged using OI arise from subthreshold synaptic and dendritic activities that parallel local field potential changes (Logothetis et al., 2001; Berens et al., 2008). It is common practice to apply a bandpass filter to the OI signal, create filtered single condition maps (SCMs) and then vector average these to obtain the classical orientation domain maps (Bonhoeffer and Grinvald, 1996; Swindale, 1998). This method has the advantage that by removing the stronger sub-threshold signals of a larger spatial scale, which are usually thought of as stimulus non-specific, one can isolate the weaker, spatially finer, signals that represent the post-synaptic spike activity (Frostig et al., 1990; Logothetis et al., 2001; Berens et al., 2008). However it has the disadvantage that, since the bulk of the haemodynamic signals arise from subthreshold synaptic and dendritic activities that parallel local field potential changes (Logothetis et al., 2001; Berens et al., 2008), any crucial information contained in signals at larger spatial scales, from which the spike output is ultimately sculpted, gets filtered out. This suggests that signals at a larger spatial scale, presumably arising from afferent axon terminals, synapses and dendrites, mainly reflect the inputs to the imaged cortical area.

With this approach, we aimed to study any orientation bias that may be present in the thalamic inputs. Anisotropies in orientation preferences have been reported in behavioural, imaging and electrophysiological studies in humans (Campbell et al., 1966; Appelle, 1972; Rovamo et al., 1982; Sasaki et al., 2006; Mannion et al., 2010; Swisher et al., 2010; Maloney et al., 2014), monkeys (Mansfield, 1974; Mansfield and Ronner, 1978; Smith et al., 1978; De Valois et al., 1982; Kennedy et al., 1985; Sasaki et al., 2006), ferrets (Chapman and Bonhoeffer, 1998; Coppola et al., 1998; Grabska-Barwinska et al., 2009) and cats (Pettigrew et al., 1968; Orban and Kennedy, 1981; Levick and Thibos, 1982; Vidyasagar and Urbas, 1982; Payne and Berman, 1983; Vidyasagar and Henry, 1990; Schall et al., 1986b; Shou and Leventhal, 1989; Li et al., 2003; Swisher et al., 2010). We investigated whether such anisotropies could be observed in the optical imaging signals we recorded from the macaque cortex. Imaging studies, especially fMRI studies in humans (Sasaki et al., 2006; Mannion et al., 2010; Swisher et al., 2010; Maloney et al., 2014) and non-human primates (Sasaki et al., 2006) have reported in the BOLD (Blood Oxygen Level Dependent) signals in V1 a ‘radial’ bias, which refers to an overrepresentation of orientations along the line joining the visual field locus to the centre of the fovea. However, fMRI signals reflect neuronal activity at a much larger spatial scale (Churchland & Sejnowski, 1988) compared to the width of orientation columns (Hubel and Wiesel, 1968) and cannot provide any information on the orientation preferences of individual neurons or orientation columns. On the other hand, with the intrinsic signals obtained from optical imaging, we can potentially analyze signals occurring at multiple spatial scales. The unfiltered optical imaging signal is capable of recording the subthreshold activity occurring at large spatial scales such as the thalamic input. The bandpass filtered images (see Fig. 1) remove them leaving only the activity at finer spatial scales, such as the responses of orientation domains, which are beyond the spatial resolution capacity of fMRI. Furthermore, studies of radial orientation bias require precise plotting of visual field loci in relation to the foveal centre and their representation on the imaged cortical surface, as we have done in this study. This is especially critical for locations close to the fovea, where errors in localising the foveal centre or receptive fields translate to greater errors in the measured radial angle than in the periphery. Thus we aimed to use OI to test two hypotheses: 1) The unfiltered, putative subthreshold signals reflecting the input will be dominated by just one or two orientations unlike the more uniform distribution of orientations seen with the filtered signals that largely reflect the spike responses. 2) The dominant orientation of the unfiltered signal will reflect the predominant orientation observed among retinal ganglion cells and in the dorsal lateral geniculate nucleus (LGN) of the thalamus, namely, the radial orientation.

**Figure 1.**
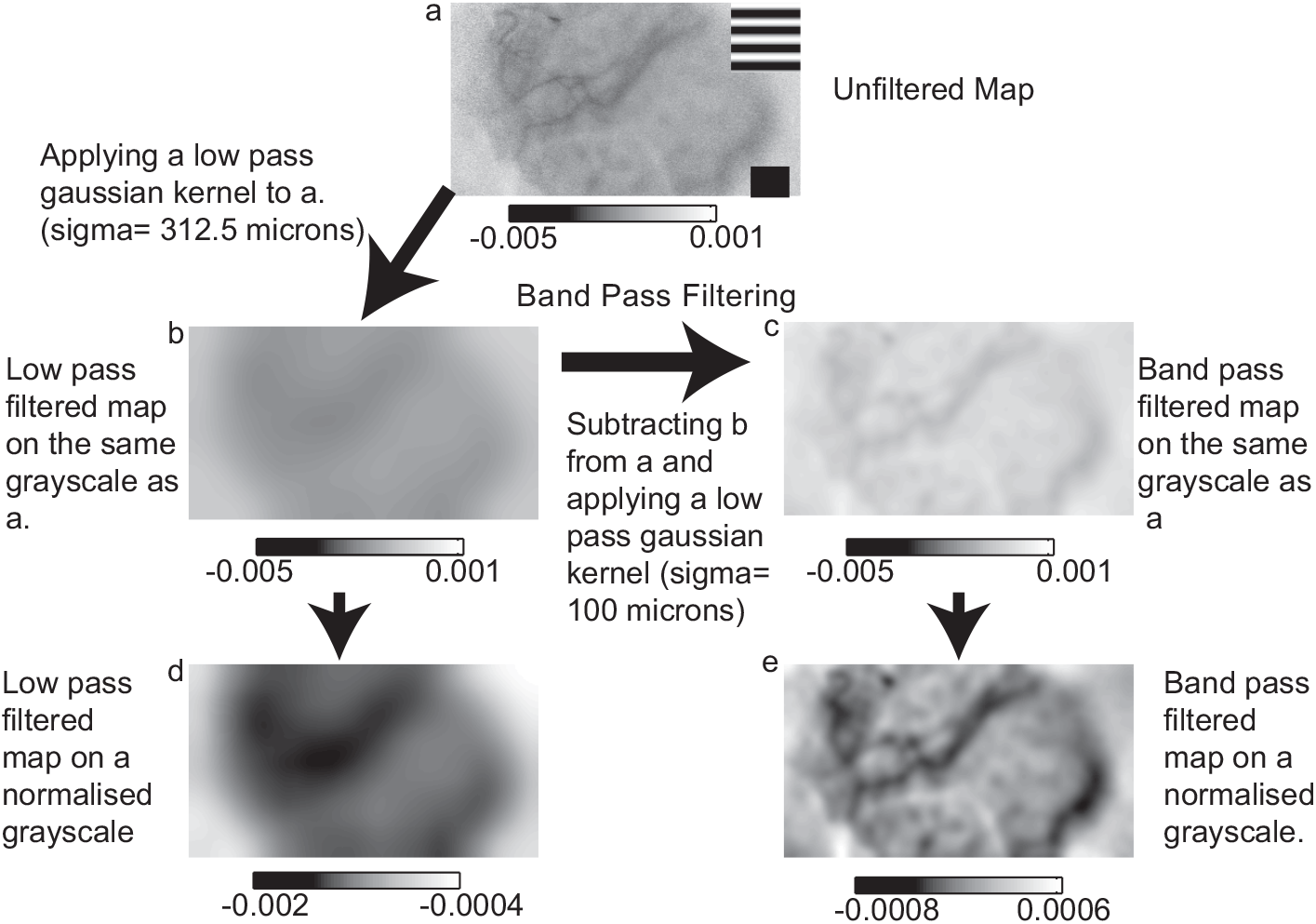
Generation of unfiltered and filtered maps from optical imaging of intrinsic signals from macaque V1. (a) An example of an unfiltered single condition map (SCM). Inset is the stimulus condition. Scale bar at bottom right is 1mm. (b) SCM after applying low pass filter (sigma= 312.5 microns). (c) Map b is subtracted from map a and a low pass filter (sigma = 100 microns) is applied to the resulting map to give the band-pass filtered map in c. Maps b and c have the same absolute grey scale as the unfiltered map in a. This shows the magnitude of the filtered signals when compared to the unfiltered signals. Maps d and e are the same as maps b and c respectively, but with their grey scales normalised to their respective minimum and maximum values, as commonly displayed in most publications to enhance contrast.

## Materials and Methods

### General and surgical procedures

Five male macaques (Macaca nemestrina; 2-4 years of age) were used in this study, which was approved by the University of Melbourne/ Florey Institute of Neuroscience and Mental Health Animal Ethics committee and conformed to the guidelines of the National Health and Medical Research Council’s Australian Code of Practice for the care and Use of Animals for Scientific Purposes. After the initial induction of anaesthesia with Ketamine (Ketamil 15mg/Kg i.m., Parnell Laboratories, Australia) and Xylazine (2 mg/kg i.m., Troy Laboratories, Australia), cephalic veins of both forelimbs were catheterized and the trachea was cannulated for artificial ventilation. Anaesthesia was maintained with Isoflurane (0.5-2%) in a mixture of nitrous oxide and oxygen (70:30) throughout the experiment. Skeletal muscle paralysis was induced with a bolus of Vecuronium (0.7 mg/kg, Organon Australia Pty Ltd) and maintained with Vecuronium (0.2 mg/kg/hr) in a mixture of 4% glucose and 0.9% saline through one of the venous catheters. End-tidal carbon-di-oxide was maintained at 3.6-3.8%. Electrocardiogram (ECG) and electroencephalogram (EEG) were recorded and used for adjusting the dosage of Isoflurane to maintain an adequate level of anaesthesia. Core body temperature was monitored using a sub-scapular thermistor, which also provided feedback to a servo-controlled blanket to maintain the body temperature at 36-37 degree Celsius. The eyes were dilated using topical administration of 0.1% Atropine (Sigma Pharmaceuticals Pty Ltd, Australia). Rigid gas permeable lenses were inserted to prevent the eyes from drying and appropriate optical lenses and 4mm artificial pupils were used to correct the refractive error and minimize optical aberrations. Craniotomy (Horsley-Clarke coordinates: 24-34 mm posterior and 2-10 mm lateral) and subsequent durotomy were performed to expose the dorsal aspect of the occipital lobe corresponding to a part of the macaque primary visual cortex (V1). A metal chamber (of diameter 10mm) was mounted over this opening using dental cement (Dentimex VA, Netherlands) and the chamber was filled with silicone oil (Poly methyl siloxane 200, Sigma Australia) and sealed tight with a cover glass.

### Stimuli

Visual stimuli were generated using a Visage stimulus generator (SDL, Cambridge Research Systems, UK) at 80 Hz on a BARCO monitor (Reference Calibrator plus; Barco Video and Communications, Belgium) kept at 57 cm from the animal. Stimuli were full-field, high contrast, square-wave gratings (1-4 cycles/deg moving at 1-1.5Hz) and presented in 8 different orientations drifting in one direction and then the other. Each grating stimulus was presented for 7.2 seconds with an interstimulus interval of 10 seconds between gratings when the animal viewed a blank screen. Data was collected for 50 presentations of each stimulus.

Stimuli for the single and multi-electrode array recordings were moving light and dark bars (length=10 degrees, width=0.2 degrees, contrast=100%, moving at 2.5 - 5 degrees per second) presented to the contralateral eye on a uniform grey screen. Multi-unit and LFP responses were collected for 18 directions (nine orientations in two directions).

### Optical Imaging of intrinsic signals

Optical Imaging of intrinsic signals (Bonhoeffer and Grinvald, 1996) was used to obtain stimulus response maps from the dorsal surface of the primary visual cortex (V1) using an imaging system (VDAQ Imager 2001, Optical Imaging, Rochester, NY). The cortical surface with its vascular details was initially imaged with an optical closed-circuit camera (Teli CS8310B, with tandem optics: 2 x Pentax lenses, f=50mm) under green illumination (545 nm). This high contrast image (green image) with the vascular landmarks (Fig. 2a) was also used as a reference image to align the response maps of the cortex. Intrinsic signals were acquired using a 630 nm red light focused at 550-700 µm below the cortical surface. The cortical region for imaging was selected to include a fairly flat surface, not only to enable good OI conditions, but also to precisely overlay the OI map on the topographical map of the visual field obtained from microelectrode recordings. For each stimulus condition, 18 data frames, each 400 ms long, were recorded. The signal-to-noise ratio was further enhanced by averaging 50 trials for each stimulus.

**Figure 2.**
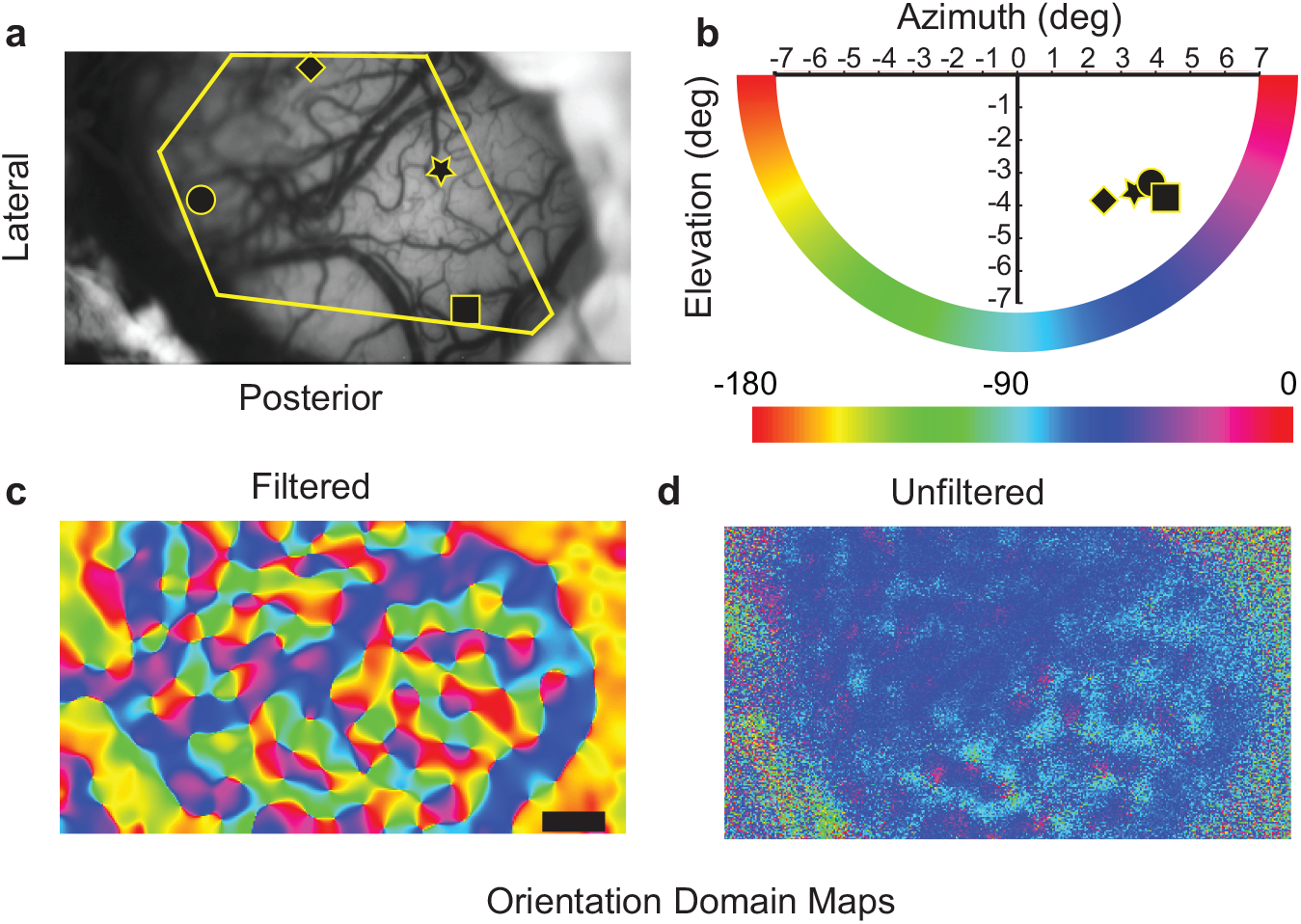
Distribution of preferred orientations in filtered and unfiltered maps. (a) V1 surface imaged under green light (λ= 545nm). The symbols indicate the locations of electrode penetrations. The yellow box indicates the cortical area used to generate the histograms in Fig. 5. (b) The receptive field locations of the electrode tracks shown in 2a. The horizontal colour scale indicates the radial angle, whose numerical value in degrees is shown. (c) Classical orientation tuning map obtained after the single condition maps were bandpass filtered as shown in Fig. 1 and vector-averaged. (d) Orientation tuning map derived from the unfiltered single condition maps. Large areas of V1 can be seen to be tuned to the same narrow band of orientations. This corresponds closely with the radial orientation of the receptive fields at the sites of reference penetrations (as in Fig. 2b). The horizontal black scale bar seen in c represents 1mm in panels a,c and d.

### Topographical recordings

Following imaging, electrophysiological recordings using lacquer coated tungsten electrodes (6-12 MOhms, FHC Inc, ME,) were performed at different locations on the imaged area. The signals were amplified (10000x; AM Systems model 1800, Washington) and filtered (300-3000 Hz for multiunit activity and in two monkeys also at 0.1-100 Hz for LFP). The signal was digitized and stored for later analysis. The receptive fields of encountered neurons were plotted on a wall chart, on which reference landmarks of the eye (optic disc parameters, vasculature and the foveal centre) were also plotted using a back projecting fundus camera. The ocular landmarks were plotted often enough to account for any ocular drifts and to enable us to obtain the precise radial angle of the recorded neurons. The receptive field locations were corrected by the fundus and foveal positions that were plotted close to the time of the single unit recordings.

### Multi-Electrode Array recordings

Multi-channel recordings were done using the TDT (Tucker-Davis Technologies, USA) system using a 16 channel linear array. The multi-electrode array (NeuroNexus Technologies Inc, USA) was inserted into V1 at an angle oblique to the surface of the cortex. The individual electrodes on the array were separated by a distance of 100 microns. The array was connected to the pre-amplifier (RA16PA, Tucker-Davis Technologies, USA) via a headstage (RA16AC). The signal was amplified (x10,000) and a band-pass filter (2.2 Hz-7.5 kHz) was applied before it was digitised (multi-unit at 12.5kHz and LFP at 1017.3 Hz) using TDT’s OpenEx software. The signal was digitally filtered between 2.2 and 100 Hz to obtain the local field potentials (LFP) and between 300 and 3000 Hz for the multi-unit recording. To obtain multi-unit responses, spikes were detected when they crossed a threshold that was 3.5 standard deviations above the noise of the respective channels.

### 1. Data Analysis

Data analysis was performed using custom scripts generated in MATLAB. Stimulus response maps were obtained as an average of 14 data frames (frames 3-16) followed by first-frame subtraction to remove illumination artefacts across 50 trials for each stimulus condition. The optimum orientation of individual pixels was calculated by vector-averaging the pixel values from the stimulus response maps (Swindale. 1998). We refer to the computed map without any image manipulation as the “unfiltered” map. The conventional orientation map was also generated by classical methods using band-pass filtering (Bonhoeffer and Grinvald, 1996; Swindale, 1998). The stimulus response maps were first filtered using a difference of Gaussian method to isolate features between sigma=312.5 um and sigma=100 um. The pixels from the resultant single condition maps were then vector-averaged to calculate their optimum orientations and provide the “filtered” maps (Fig. 1).

We used the receptive field locations in reference penetrations, projections of the foveal centre using the fundus camera and cortical magnification factor values across the retina (Dow et al., 1981) to determine the radial angle of points on the cortical surface that were uniformly spaced 375 µm apart (Fig. 3). We then calculated the average orientation response of pixels in a 750 µm square (‘optimum orientation’) around each of the points for both the filtered and unfiltered conditions. We then compared the computed radial angle and the optimum orientation to determine to what extent the OI response preferred stimuli close to the radial orientation. We also vector-averaged the response of every pixel and collected the data from all 5 animals in one histogram.

**Figure 3.**
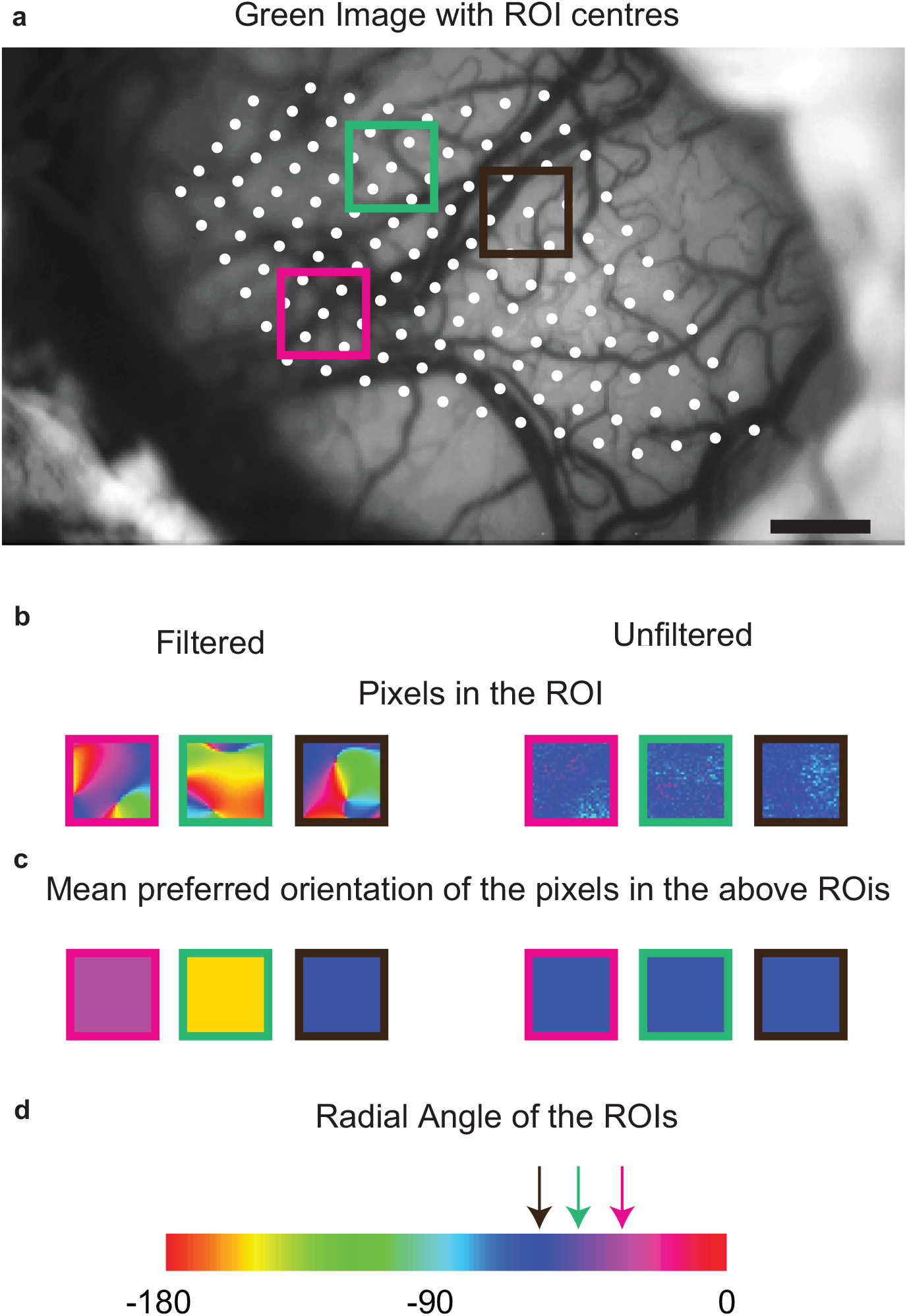
Construction of ROIs for analysis. (a) Location of the centres (white dots) of the regions of interest (ROIs) are shown on the green image of one of the imaged cortices. Iso-azimuthal and iso-elevation lines were constructed from the electrode tracks and their respective receptive field locations (see Materials and Methods). Then, points 15 pixels apart were marked along these lines to denote the ROI centres and 30 x 30 pixel ROIs were placed around the centres (coloured boxes show 3 examples). The size and distance between the ROIs were so chosen to avoid sampling within a single orientation column repeatedly. (b) shows the three coloured boxes of a and the pixels within the ROI for the filtered and the unfiltered maps. The average vector of the pixels within the ROIs are shown in (c). The radial orientation of the ROI was calculated using the azimuth and elevation of the centres and the optimum orientation was obtained by calculating the circular mean of the pixels within the ROI in the vector averaged map. (d) shows the colour scale used to generate the orientation domain maps. The radial angles of the three coloured boxes in (a), (b) and (c) are shown by the arrows above the colour scale.

## Single and Multi-electrode array recordings

The spikes were collected in 20 ms bins to create peri-stimulus time histograms (PSTHs) and smoothed across three bins to generate the spike density function (SDF) from 10 trials for each orientation. Response in the peak bin in the SDF was taken as the response for the particular orientation. To obtain LFP responses to bars, the recorded signal was filtered between 20 and 70 Hz to reveal only the activity in the gamma frequency range. The magnitude of the response was then calculated as the average of the maximum peak-to-trough amplitudes in the responses in individual trials.

## Results

We used optical imaging of intrinsic signals to image an exposed area of the primary visual cortex in five anaesthetised macaques (in three the left hemisphere and in the other two the right). The haemodynamic responses of the imaged area in response to gratings of 8 different orientations (0-157.5 degrees) were recorded. A single condition map (SCM) was generated for response corresponding to each orientation using the first frame subtraction method (see Materials and Methods). These SCMs were then used to produce either the unfiltered SCM (as in Fig. 1a) or the filtered SCMs (as in Fig. 1e). The unfiltered and the filtered SCMs corresponding to each orientation were vector averaged (Swindale, 1998) to produce the unfiltered and filtered orientation maps respectively.

When no band-pass spatial filter was applied to the haemodynamic signal and an unfiltered orientation map was created, we found that large areas of the cortex were tuned to only a narrow range of orientations (Fig. 2d), but this was not the case for the filtered map (Fig. 2c). To precisely ascertain whether the dominant orientation observed in the unfiltered map bore any relationship to the radial orientation, we made multi-unit recordings from microelectrode penetrations made perpendicular to the cortical surface (Fig. 2a) and obtained a map of receptive field locations within the imaged area, using receptive field locations of the reference penetrations and published values of cortical magnification factor for the macaque (Dow et al., 1981). The projection of these recorded sites on to the visual field (Fig. 2b) reveals the radial orientation of these cortical sites. There is a close correspondence between these radial orientations and the narrow range of dominant orientations in the unfiltered map (refer to the pseudocolour key between Fig. 2b and Fig. 2d). Results of similar correspondence in the other four animals are shown in Supplementary Fig. S1. As in Fig. 2, the radial angle is seen to be overrepresented in these unfiltered maps.

## Orientation Preferences of Regions of Interest

In order to quantify any preference for the radial orientation in the cortical OI maps, analysis was done on a large number of small regions of interest (ROIs) within the imaged area of each of the five animals (see Fig. 3). For 750 × 750 µm areas (30x30 pixels) spaced 375 µm apart, the radial orientation was calculated (as described above and in Materials and Methods). The preferred orientation of the signal was calculated from the OI data by taking the mean vector of the individual pixels within an ROI. This was done separately for the unfiltered and filtered orientation domain maps. The absolute difference between the radial orientation of the visual field locus of the ROI and optimum orientation of the intrinsic signal of the ROI was calculated for both the filtered and unfiltered conditions for all 5 animals (n=456). These are shown as a histogram in Fig. 4a. The distribution for the unfiltered and filtered maps were significantly different (χ^2^=283.01; df=3; p<0.0001). For the unfiltered maps, the majority of ROIs were tuned to the radial orientation and the distribution was significantly different from a uniform distribution (χ^2^=505.28; df=3; p<0.0001). For the filtered maps, although the distribution of absolute differences was significantly different from a uniform distribution (χ^2^=35.21; df=3; p<0.0001), the preference for the radial orientation was markedly less than in the unfiltered maps as can be seen in Fig. 4a.

**Figure 4.**
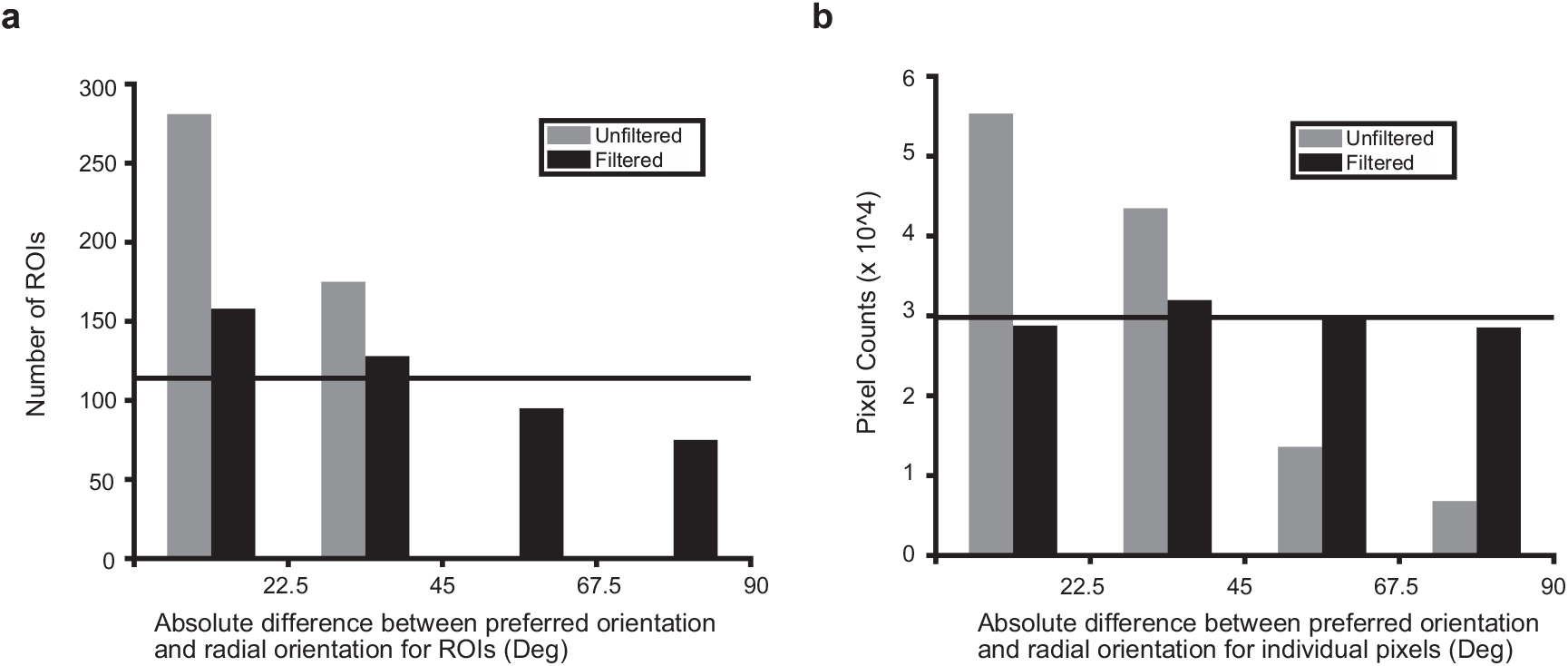
Relationship between the preferred stimulus orientation of the intrinsic signal at a site on the V1 cortex and the radial angle of the site in the visual field. (a) Distribution of absolute difference between the radial angle of each ROI (as defined in Materials and Methods) and the angle of the vector-averaged OI response of the unfiltered and filtered maps. The horizontal line indicates the expected number of ROIs in each bin if the distribution were uniform. (b) Distribution of the difference between preferred orientation and the radial orientation of individual pixels in the unfiltered and filtered maps. The optimum orientation of single pixels was subtracted from the mean radial angle of the whole analysed area. The subtracted values were then grouped in 22.5 degree bins.

## Orientation Preferences of Single Pixels

The analysis was repeated for individual pixels (n=119229) instead of just for the (30x30 pixel) ROIs. The results (Fig. 4b) were similar. The distributions of the unfiltered and filtered maps were significantly different (χ^2^=54077; df=3, p<0.0001). For the unfiltered maps, the majority of pixels were tuned to the radial orientation and the distribution was significantly different from a uniform distribution (χ^2^=54691; df=3, p<0.0001). For the filtered maps, the distribution of absolute differences was also significantly different from a uniform distribution (χ^2^=246.24; df=3, p<0.0001), but there is no preference for the radial orientation.

However, with this analysis using single pixel values (Fig. 4b), the very large sample sizes can potentially lead to statistically significant differences from a uniform distribution even when this difference is small. However, as per our hypothesis, the deviation from a uniform distribution of orientations should be much stronger for the unfiltered samples than for the filtered. Though this difference can be seen in the histograms shown in Fig. 4, in order to control for the effect of sample size, we performed a bootstrapping exercise using fewer numbers of randomly sampled pixels. We sampled 1000 times (i.e., 1000 trials) with replacement from the original data sets of the unfiltered and filtered maps. We set two sample sizes, 40 pixels per trial (Fig. 5) and 1000 pixels per trial (Fig. 6). Box plots in a and b show the distribution of the data points (pixels) with respect to the radial orientation (set as 0 degrees) for each trial in the filtered and the unfiltered conditions respectively. The distribution of the chi-square test statistic obtained for the 1000 trials is shown in panels c and d for the filtered and unfiltered conditions respectively. When the sample size was set to 40 (Fig. 5c), the individual pixels in the filtered condition were indeed uniformly distributed (Mean χ^2^ over 1000 trials = 2.99, 95% confidence interval = [2.84, 3.13]; the critical value for the 3 degrees of freedom being 7.81 for p=0.05) but the individual pixels in the unfiltered condition (Fig. 5d) were significantly different from a uniform distribution (Mean χ^2^ over 1000 trials =21.09, 95% confidence interval = [20.95, 21.24]). When the sample size was set to 1000, the same pattern was observed for the pixels sampled from the filtered maps (Fig. 6c; Mean χ^2^ over 1000 trials = 5.18, 95% confidence interval = [4.92, 5.44]) as well as the unfiltered maps (Fig. 6d; Mean χ^2^ over 1000 trials =461.93, 95% confidence interval = [461.68, 462.19]). This suggests that the statistically significant result obtained for the difference between the radial orientation and optimum orientation of single pixels was largely an artifact of sample size.

**Figure 5.**
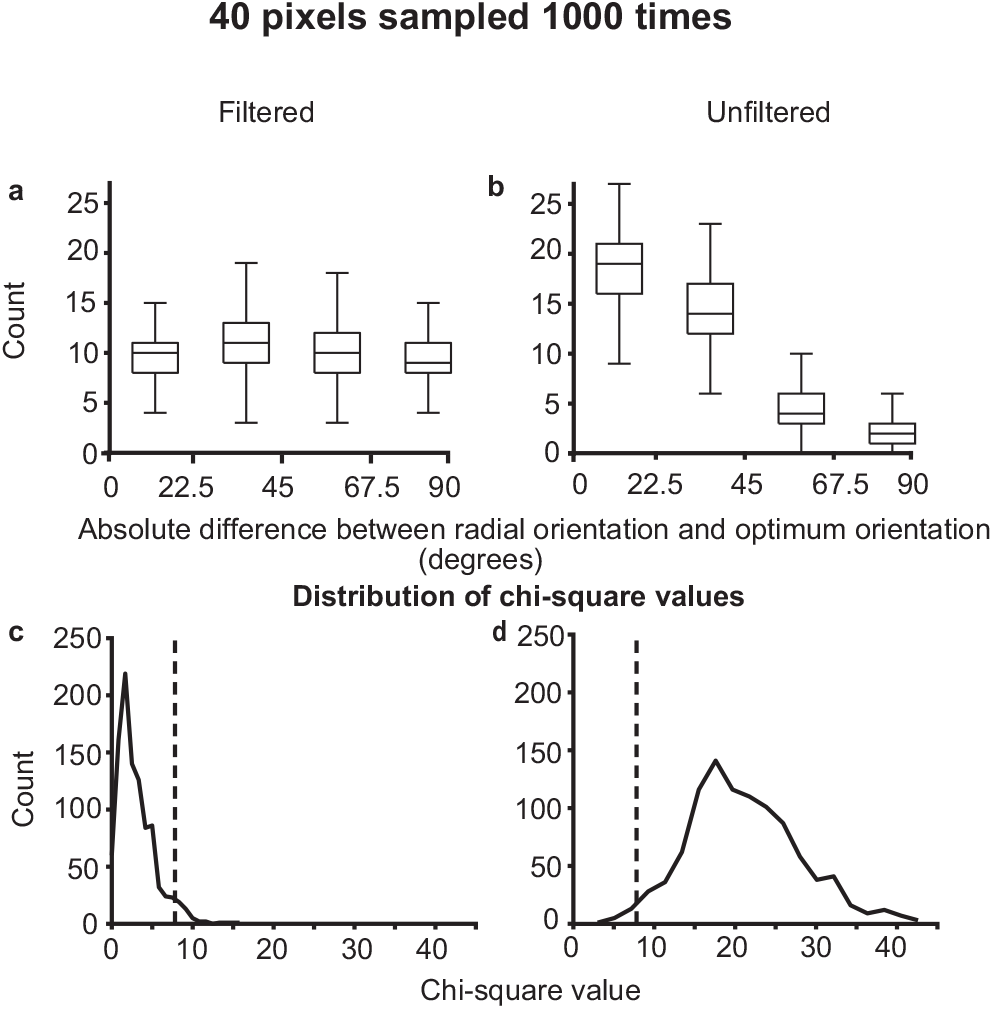
Distribution of preferred orientations for subsets of 40 single pixels for filtered (a) and unfiltered (b) data sets in relation to the radial orientation of the imaged area, with zero representing no difference. The distribution in each trial was compared to a uniform distribution and the χ^2^ values for the 1000 trials are shown for the filtered (c) and unfiltered (d) data sets. The vertical dotted line in c and d indicate the critical χ^2^ value (7.81) for df=3 and p<0.05.

**Figure 6.**
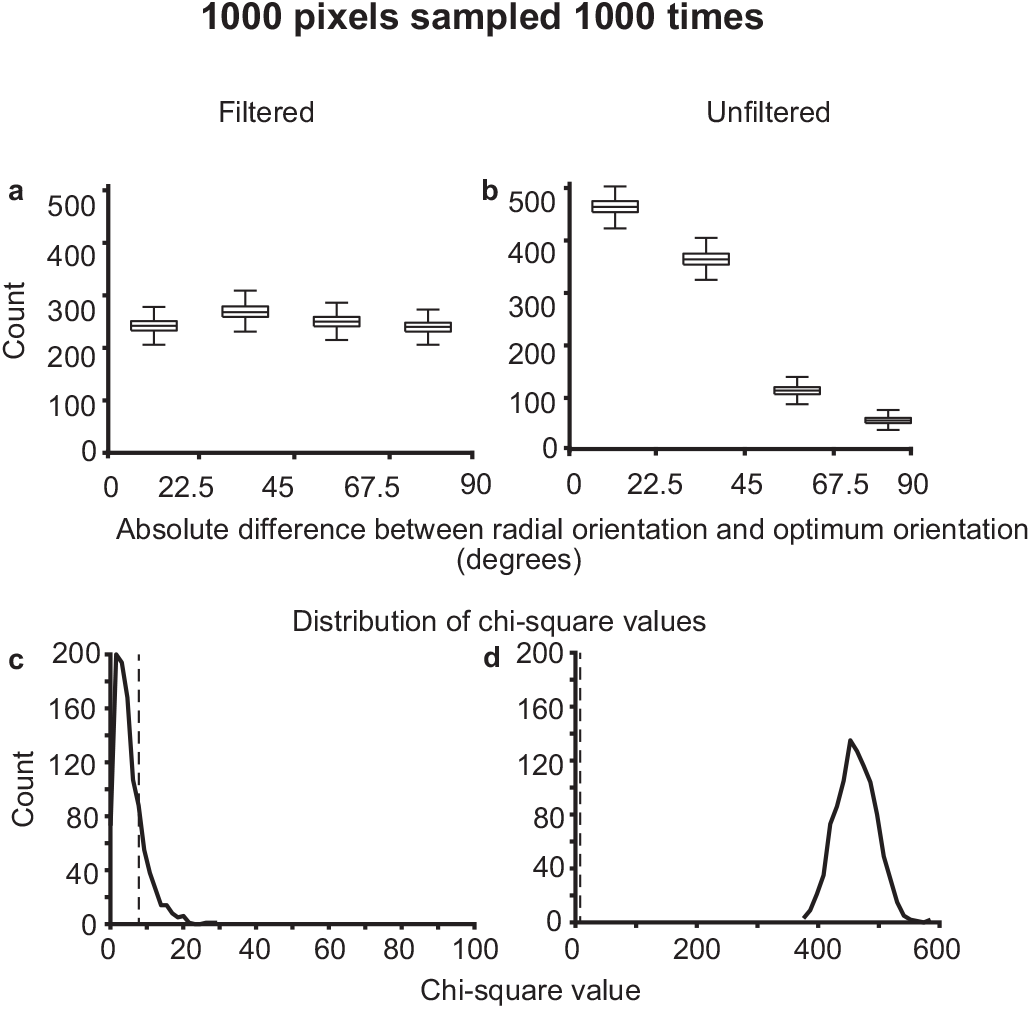
Distribution of preferred orientations for subsets of 1000 single pixels. See figure legend for Fig 5 for full description.

## Comparison with Local Field Potential data

In two animals, we recorded LFP and multi unit responses from six locations using single tungsten microelectrodes and in one animal, we used a 16-channel multi-electrode linear array, placed in the cortex at an angle oblique to the surface in the imaged area, but mostly within the supragranular layers. Multi-unit activity (MUA) and local field potentials (LFP) were recorded in response to bars of different orientations. The orientation tuning curves of the MUA and LFP responses are shown for the multi-electrode array in Figs. 7a and 7b respectively. The multi-unit activity was fairly well tuned to orientation and showed considerable variation from electrode to electrode, whereas the LFP response showed broader tuning and much less variation in its preferred orientation. The poor correlation between the preferred orientations of the MUA and LFP was similar to what was seen in an earlier study on the macaque (Berens et al., 2008). However, we also found that the orientation of the LFP response was within 22.5 degrees of the radial angle in half the cases (11/22), with the distribution of differences significantly different from a uniform distribution (n=22; χ^2^=8.18; df=3, p=0.04), whereas the orientation of the multi-unit response was more evenly distributed (n=22; χ^2^=3.09; df=3, p=0.38). These results are consistent with the data presented in Fig. 4, where the unfiltered signal mostly preferred the radial orientation.

**Figure 7.**
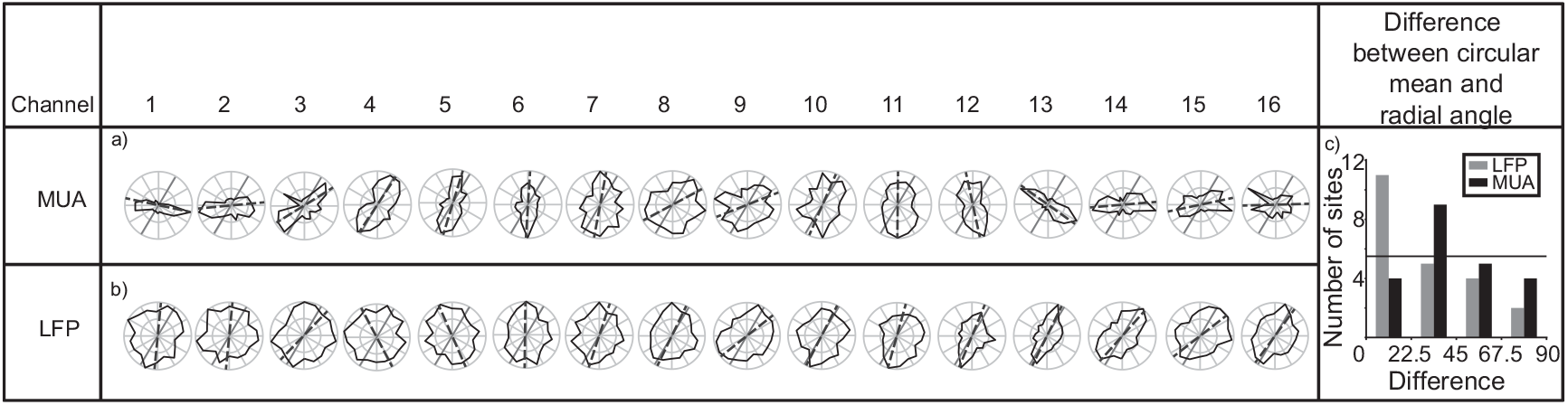
Multi-unit activity (MUA) and local field potential (LFP) responses at each site of the 16 channel linear array placed in the imaged area. Orientation tuning curves of the multi-unit (a) and the LFP (b) responses to a light bar are shown, normalised to the maximum response at each channel. The continuous gray line indicates the average radial angle of the imaged area and the dashed black line is the circular mean of the multi-unit recording in (a) and the LFP response in (b). (c) shows a histogram of the differences between the circular mean and the radial angle of the multiunit response (black bars) and the circular mean and the radial angle of the LFP response (grey bars) for the 16 channels shown in (a) and (b) along with the LFP and response measured from the six single electrode recordings. Horizontal line indicates the expected value for a uniform distribution.

## Discussion

Our optical imaging and electrophysiological results both suggest that the presynaptic, synaptic and dendritic activity occurring over a broad spatial scale covering many classical orientation domains are dominated by a bias to one stimulus orientation, namely the radial, whereas the orientation domains themselves, consisting of clusters of cells known to exhibit sharp orientation selectivity in their spike responses, show a reasonably full range of orientation preferences. We believe that the orientation selectivity and the radial bias observed in the unfliltered maps reflect the biases in the thalamic inputs to the cortex and the subthreshold activity within the cortex. If most thalamic inputs to cortex are indeed tuned to the radial orientation, it has important implications for the development of cortical architecture. It is consistent with the proposal that the signals conveying information on stimulus orientation arrives in the cortex in a small number of broadly tuned channels and that the cortex develops the whole range of orientation selectivities from these inputs, similar to the well-known instance of primate colour vision (Vidyasagar, 1985; Vidyasagar and Eysel, 2015). Such a mechanism giving rise to a gradual change in orientation preference in the form of the classical orientation domain architecture (Hubel and Wiesel, 1968; Vidyasagar and Eysel, 2015) is comparable to the organisation demonstrated for other cortical features, especially for ocular dominance (Hubel and Wiesel, 1968) and ON/OFF domains (Kremkow et al., 2016). Though there is no significant trend for a second preferred orientation in our unfiltered optical imaging maps, we argue that this may not be necessary. Since we know that in colour vision, not only are the short-wavelength cones much sparser than the other two cones, but even the ratio of L and M cones vary widely between individuals with trichromatic vision, who all have, nevertheless, very similar colour vision (Kremers et al., 2000). Presumably, at the cortical level, the weighting between the different channels are adjusted to compensate for the difference in the numbers of different cone types and finally provide the full range of trichromatic colour vision. We propose that a similar process occurs in the orientation domain, even though the actual numbers of afferents preferring non-radial orientations may be much fewer than for the radial.

It should be noted that we focused the camera at 550-700 microns below the cortical surface, which is just above the interface between layer 3 and the thalamic input layer 4. The majority of layer 4 cells in the macaque V1, except those in the sublayer 4B, show poor orientation selectivity, largely reflecting the responses seen in the geniculate. The first major sites of large-scale generation of sharp orientation selectivity then are in fact the supragranular layers, 2 and 3 (Bullier and Henry, 1980; Hawken and Parker, 1984; Hubel and Wiesel,1968). Furthermore, macaque supragranular layers receive a direct koniocellular input from the geniculate (Klein et al., 2016), which again would reflect the broader, subcortical orientation preferences. Hence, even though we focused the camera at the bottom of layer 2/3, one can conclude that the orientation of the larger spatial scale inputs reflects the orientation biases of the thalamic inputs.

Our study also establishes the value of observing the OI signal at different spatial scales to reveal underlying processes as has been pointed out in an fMRI study done on the primary visual cortex of cats and humans (Swisher et al., 2010) and as recently demonstrated also for macaque area V4 (Tanigawa et al., 2016).

Furthermore, our findings resolve an apparent discrepancy in the earlier literature about orientation anisotropies in the visual system. Studies that either recorded single unit activity or mapped orientation domains using optical imaging of intrinsic signals have reported a preponderance of neuronal activity preferring cardinal (i.e., horizontal and vertical) orientations in the primary visual cortex of cats (Pettigrew et al., 1968; Orban and Kennedy, 1981; Payne and Berman. 1983; Vidyasagar and Henry, 1990), monkeys (Mansfield, 1974; Mansfield and Ronner, 1978; De Valois et al., 1982; Kennedy et al., 1985; Smith et al., 1990) and ferrets (Chapman et al., 1998; Coppola et al., 1998; Grabska-Barwinska et al., 2009). This was generally consistent with the classical oblique effect in humans (Campbell et al.,1966; Appelle, 1972; Rovamo et al.,1982). However, as noted earlier, fMRI studies of V1 in humans (Sasaki et al., 2006; Mannion et al., 2010; Swisher et al., 2010; Maloney et al., 2014) and macaques (Sasaki et al., 2006) have generally shown a radial bias.

Against this backdrop of controversy, it is worthwhile to point out that every one of the above studies is confounded by one or more of the following three factors: 1) Human psychophysical studies measure the overall response of the visual system and tell us little about the orientation anisotropies that may or may not exist at earlier stages. It has been claimed that the oblique effect may be a phenomenon happening more centrally along the visual pathway rather than at the level of orientation detectors of area V1 (Westheimer, 2003). 2) Without ascertaining the precise relationship of each receptive field to the fovea, electrophysiological recordings of single cells from V1 may be able to inform us only about any preponderance of cardinal versus oblique orientations, but not of radial orientations, as they will fail to reveal the radial angle of the receptive field locations relative to the fovea, especially when they are closer to the foveal centre. Any radial bias will also manifest as cardinal preferences if sampling were biased towards vertical or horizontal meridians, since the visual field representations of recording sites are usually constrained by technical accessibility. 3) Human neuroimaging studies lack the spatial resolution that is necessary for imaging at the level of single neurones or even single orientation domains. Thus, whether the radial biases seen extensively at retinal and LGN levels initially dominate the input to the cortex but get transformed by cortical circuitry to a more uniform representation in the spike outputs of the typical cortical cells showing sharp orientation selectivity, has in fact been an open question. Our results resolve these confounds and show the more uniform distribution of preferred orientations seen in orientation domains and single cells emerging from an input that is dominated by a radial bias.

In one of our previous studies on cats (Vidyasagar et al., 2015), we reported that the thalamic inputs tended to have the same orientation preferences as the orientation columns where they terminated in, but with a notable degree of scatter in the relationship (Pearson r = 0.63). Those results are indeed compatible with our claim here that the majority of the inputs to the cortex were tuned to the radial orientation. Apart from a possible species difference, the fair degree of scatter around the identity line in that study is a measure of the departure from the radial angle of ROIs and pixel vectors in the filtered maps of the present study.

While the bias for radial orientation may serve the function of detecting optic flow, especially in peripheral vision (Sherk et al., 1997), it has been suggested that the growth of the retina from centre outwards may inevitably lead to the majority of retinal dendritic fields being radially elongated, underlying the bias for radial orientations in the visual system (Schall et al., 1986a, 1986b). We suggest that this developmental constraint is being exploited by the visual cortex to encode a range of orientations by combining the outputs of these neurons with those that are tuned to non-radial orientations, though the latter may be smaller in relative numbers. Thus, as in colour processing, orientation discrimination may also originate in the retina - in the broad selectivities shown by retinal cells around only a limited number of peak orientations (Vidyasagar and Eysel, 2015).

## Acknowledgments.

The work was supported by an Australian National Health and Medical Research Council Project Grant (APP1060610) to T.R.V.

